# Social experiences shape song preference learning independent of developmental exposure to song

**DOI:** 10.1101/2023.05.11.540402

**Authors:** Erin Wall, Sarah Woolley

## Abstract

Communication governs the formation and maintenance of social relationships. The interpretation of communication signals depends not only on the signal’s content, but also on a receiver’s individual experience. Experiences throughout life may interact to affect behavioral plasticity, such that a lack of developmental sensory exposure could constrain adult learning, while salient adult social experiences could remedy developmental deficits. We investigated how experiences impact the formation and direction of female auditory preferences in the zebra finch. Zebra finches form long-lasting pair bonds and females learn preferences for their mate’s vocalizations. We found that after two weeks of cohabitation with a male, females formed pair-bonds and learned to prefer their mate’s song regardless of whether they were reared with (“normally-reared”) or without (“song-naïve”) developmental exposure to song. In contrast, females that heard but did not physically interact with a male did not prefer his song. Moreover, even following mating, song-naive females failed to show species-typical preferences for courtship song, hinting that mating is insufficient to ameliorate broader deficits in sensory processing that arise during development. Thus, courtship and mating interactions, but not acoustic-only interactions, strongly influence preference learning regardless of rearing experience, and may dynamically drive auditory plasticity for recognition and preference.

## INTRODUCTION

Sensory systems are selectively adapted to prevailing environmental conditions, not only through evolution but also over the course of development [1–3]. Indeed, across auditory, visual, olfactory, somatosensory, and vestibular systems, the local environment during development shapes neuronal responses and these changes in the sensory perception of the external world can shape physiology and behavior throughout the lifespan [4]. For example, in the paradigm case of developmental plasticity, the visual system, monocular deprivation early in life leads to persistent changes in visual acuity through adulthood [5]. Similarly, the auditory cortex is highly plastic in young animals. In juveniles, auditory experiences that promote the progressive development of tonotopic organization and frequency-selectivity or tuning are key to the structural organization and signal processing capacities of the primary auditory cortex and associated auditory perceptual abilities throughout life [6,7].

Experience can also shape sensory responses in adults. In particular, while developmental sensory experiences and manipulations can have lasting effects on sensory organization, social experiences can lead to sensory learning and plasticity in adults even after the closure of developmental sensitive periods. In a number of species, mating can lead to learning and plasticity, especially in species in which mating induces the formation of pair bonds: selective, enduring, and often monogamous relationships [8–11]. Pair bonded individuals recognize and prefer their mate or their mate’s signals (vocalizations, scent, etc) over unfamiliar individuals, indicating that there is plasticity and learning of sensory signals or representations of a mate in bonded animals [12–15]. For example, both male and female prairie voles show recognition of and preference for their mate’s chemical signature [16,17]. Similarly, female zebra finches prefer the song of a mate over the song of an unfamiliar conspecific, and this preference is accompanied by a habituation of immediate early gene expression in the secondary auditory cortex that is similar to the habituation seen to songs of other familiar individuals, such as the father or tutor [18–21]. However, while adult social experiences can dramatically impact auditory plasticity and perception, developmental organization during a critical period may contribute to or limit the impact of adult experience. Further, the degree to which developmental auditory exposure may be necessary for animals to learn to recognize a mate’s vocalizations, or whether adult experiences can ameliorate developmental deficits in auditory processing, is unclear.

Songbirds offer a unique opportunity to study how interactions between developmental and adult auditory experience shape auditory learning and recognition. In zebra finches, females use the learned songs of males to recognize individuals and are sensitive to subtle changes in song indicative of the male’s social and motivational state [21–23].

Birds reared without exposure to song (‘song-naïve’ birds) differ from normally-reared birds in their ability to discriminate frequency, their behavioral preferences for species-typical courtship songs, and in the neural activity driven by particular songs [22,24–26]. Finally, adult social experiences such as mating can drive learning and plasticity of female song preferences, potentially through activation of mesolimbic dopamine regions [18,21,27–30]. Here, we investigated the effects of mating experience, as compared to listening to a male’s vocalizations, on preference learning in adult birds, and whether females can learn to prefer a mate’s song even if they lack developmental song exposure. Compared to females reared with song exposure (‘normally-reared’), we find that females raised without song exposure (‘song-naive’) show similar pair-bonding behaviors with a male and preference learning for a mate’s song, even though the mate is the first male they have encountered. However, mating is not sufficient to promote consistent species-typical preferences for courtship songs like those seen in normally-reared birds. Interestingly, this plasticity of preference is only observed in females whose social experience with a male was multimodal rather than acoustic-only.

Females allowed to hear, see, and physically interact with a male learned to prefer his song, while females that only listened to the male did not. Together, these data highlight that while plasticity and mate preference learning are maintained even in the absence of developmental auditory exposure critical for shaping auditory organization, there are lasting changes to auditory processing that are not rescued by adult social experiences.

## METHODS

### Animals

All zebra finches (n=86 females and 24 males, average 9 months old, range 3-26 months) were maintained on a 14:10 light:dark cycle, given ad libitum access to seed, grit, and water, and weekly supplements (e.g. lettuce, egg). We raised females in one of two conditions. One set of females (‘normally-reared’) was raised to 60 days post hatch in a cage with both parents and siblings. A second set of females (‘song-naïve’) were raised in sound-attenuating chambers (‘soundboxes’; TRA Acoustics, Cornwall, Ontario) with only the mother and siblings. Specifically, fathers were removed within five to seven days post-hatch, prior to the period when females memorize the father’s song [31] and male siblings were removed at 30-40 days post-hatch, prior to producing stereotyped song [32]. After 60 days, normally-reared-females (n=48) were housed in same-sex group cages in our mixed-sex colony, and song-naïve females (n=38) were housed in same-sex group cages in an all-female colony. All males used for social cohabitation or song recordings were normally-reared and housed in same-sex group cages in the mixed-sex colony prior to use. All procedures adhered to Canadian Council on Animal Care guidelines and the protocol approved by the Animal Care Committee of McGill University.

### Cohabitation conditions

To assess the degree to which female preference is shaped by specific social experiences in adulthood, females were either housed in a cage with a male (‘mated females’) or housed in a cage with another female (‘unmated females’) (Figure 1A). The cages containing mated and unmated females were both located within the same soundbox but separated from each other by an opaque barrier. This meant that all birds could hear each other, but only see and physically interact with their cage mate. Both cages were provided with nesting material weekly and remained together for two weeks.

**Figure 1.**
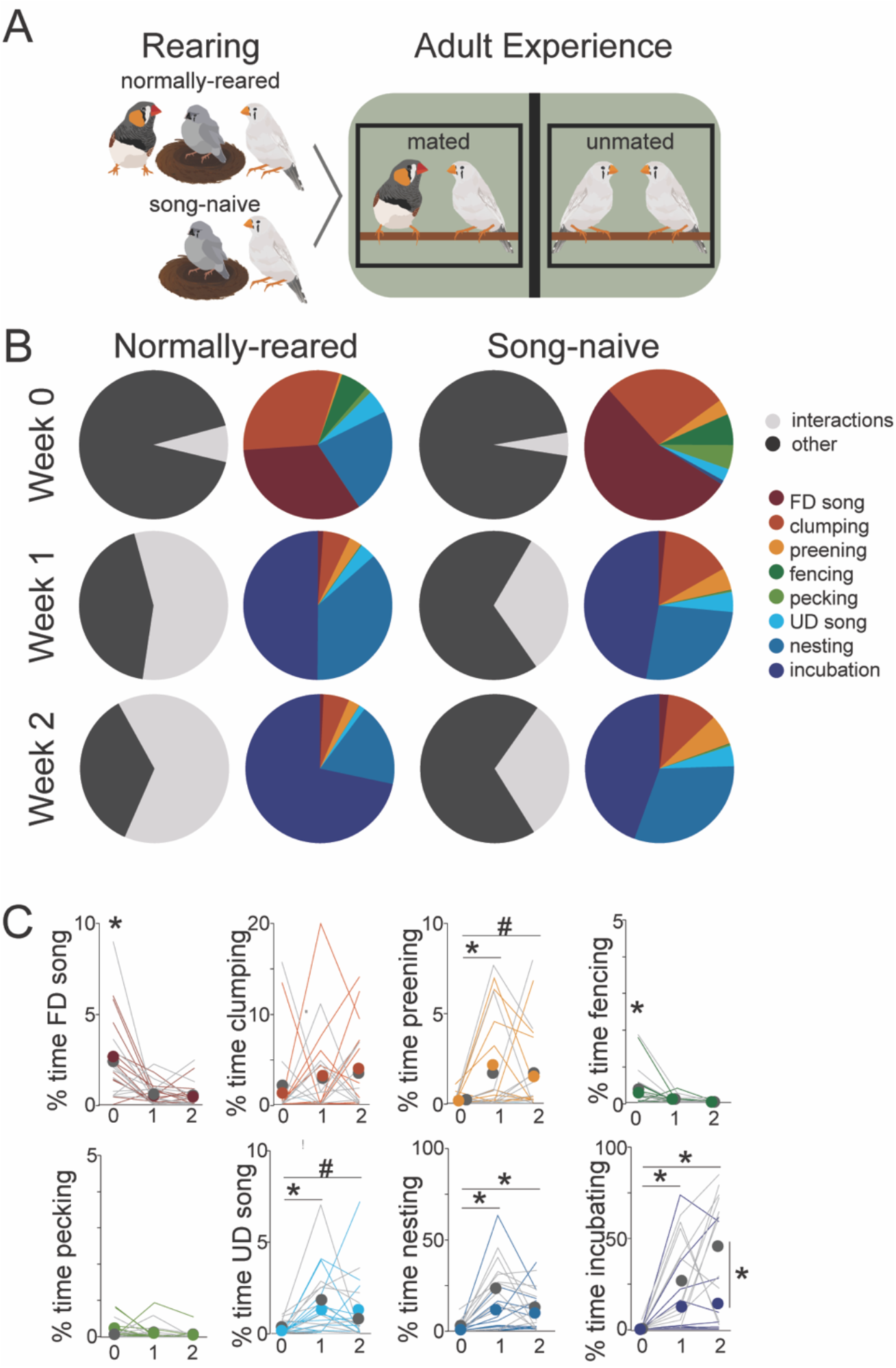
Normally-reared and song-naïve females exhibit similar progressions of pair bonding behaviors over two weeks of cohabitation. A) Experimental set-up. Females were reared either with both parents (‘normally-reared’), or without a father (‘song-naive’; male - orange cheeks, juvenile female – in nest, adult female - light grey). As adults, normally-reared and song-naive females were housed in pairs, with either a male (‘mated’, left) or a female (‘unmated’, right), in the same sound-attenuating chamber (green) separated by an opaque barrier (black). Pairs were filmed for 2-hours immediately after the first introduction (Week 0), and after one (Week 1) and two (Week 2) weeks of cohabitation. (B) Mated, normally-reared and song-naive females show overall similarities at each time point in the total percent of time spent interacting (left) and the time spent interacting broken down into component behaviors (right). See Methods. C) Data for all individuals (lines) and means (points) for the percent of the 2hr video recording spent engaging in interactive behaviors by normally-reared (gray) and song-naïve (color) females. Significant differences over time are indicated above, significant differences between rearing conditions are indicated vertically. *p<0.05, #p<0.10.

### Video recording

We filmed each group when the male was first introduced (‘Week 0’), and after the first (‘Week 1’) and second (‘Week 2’) week of cohabitation using a GoPro Hero or Hero 5 camera. All recordings lasted for at least 2 hours and were recorded at the same time of day within each group. In a few cases (n=18) we recorded birds continuously over 2 weeks. For these birds, we analyzed 2h of video from the three time points listed above.

### Preference testing

To assess preferences for male songs, females were given a two-choice active choice task as described previously [33]. Briefly, females were placed in a cage containing two strings that, when pulled, would each trigger the playback of song from a single male zebra finch through an adjacent speaker (e.g., song of male A for one string and song of male B for the other string). Females were trained to associate pulling strings with sound playback using either conspecific and heterospecific songs or calls from unfamiliar females. Each test consisted of two two-hour sessions. To control for side bias, contingencies switched for the second session of every test. For both sessions, the session started once a female pulled each string three times. Each female was tested on at least three stimulus sets: familiar male vs. unfamiliar male, courtship vs. non-courtship songs from a familiar male, and courtship vs. non-courtship songs from an unfamiliar male. The order of stimulus presentation was randomized within and across testing days for each female. Females were tested on 1-2 stimulus sets per day for a maximum of five consecutive days, including acclimation time. If females did not complete the necessary tests after five days, they were housed in small same-sex groups for at least 2 days before resuming testing. We found that these brief pauses in testing improved motivation to hear song and complete the tests.

### Song Stimuli

Songs were recorded as previously described [21,22,30]. Briefly, males unrelated to the experimental subjects were recorded in sound attenuating chambers with omnidirectional microphones (Countryman Associates, California) using a sound-activated recordings system (Sound Analysis Pro, SAP; 44.1kHz)[34]. Unfamiliar females not involved in the experiment were presented to the male in a separate cage for brief intervals to collect female-directed song. Songs selected for stimuli were a representative sample of the male’s typical variation in song duration, number of motifs, and introductory notes. All songs were free of noise and female calls. Each stimulus set consisted of 8-15 songs from each male. When females were tested on the songs of two different males, males with similar durations and numbers of motifs were selected as stimulus sets. For familiar male vs. unfamiliar male tests, we used a yoked design: each male’s song served as the familiar song for one set of females (one mated and two unmated, see Cohabitation conditions), and the unfamiliar song for a different set of females. Therefore, the same stimulus set is used for two sets of females whose experience with the two males differ. All stimulus songs were bandpass filtered (300–10 kHz), normalized by their maximum amplitude, and saved as wav files (44.1 kHz) using custom written code in Matlab (Mathworks, Natick, MA).

### Analyses

#### Pair Behaviors

The three 2h samples of video taken at Weeks 0, 1, and 2, were quantified by individuals blind to the rearing condition of the females. We recorded the duration (in seconds) and number of occurrences of a number of individual and interactive behaviors: courtship singing, non-courtship singing, clumping, allopreening, pecking, bill fencing, and nesting behaviors [30,35–38]. Courtship singing by the male was defined by a display of at least two of the following behaviors during singing: orienting towards the female, fluffing of body feathers while flattening feathers on the head, and courtship dancing [21,30,38,39]. We calculated the percent of time spent on each behavior out of two hours for each pair.

### Preference tests

From each test, the total number of playbacks for Stimulus A and Stimulus B during each session is determined. In addition to quantifying the number of pulls for each stimulus, we also performed a bootstrap with replacement (10,000 iterations) to attain a normalized metric of preference strength (‘preference index’) with 95% confidence intervals (CIs). Females were considered to have a significant preference if the mean and 95% CIs were above or below the chance threshold (0.5). We used mixed-effects models with bootstrapped preferences as the dependent variable and experience, rearing, and the interaction of experience and rearing as independent variables and individual ID as a random variable. For each rearing and experience condition, we also tested whether the distribution was significantly different from chance (0.5). Finally, we also coded responses as either a significant preference or no preference based on the bootstrapped preference and CIs. Preferences were coded as 1 (above 0.5) or -1 (below 0.5) if the CIs did not cross 0.5, and as 0 if the CIs did cross 0.5. We analyzed variation in the pattern of significant preferences across groups using nominal logistic models with rearing, experience, and their interaction as independent variables.

We investigated the correlation between preference strength and individual pair bonding behaviors in two ways. First, we performed a correlation analysis for each rearing condition and time point between the preference score and each of the pair bonding behaviors. To gain a greater sense of how combinations of behaviors relate to preference, we also used a principal components analysis (PCA) to investigate the correlations between pair bonding behaviors and mate preference. We ran separate PCAs for each time point (Week 0, 1, 2) and rearing condition and included all pair-bonding behaviors and the bootstrapped preference score. We selected the component (in all cases, the first or second PC), with the highest loading of preference. We then compared the loading matrices for each of these components across time and rearing condition. All statistical analyses were completed using JMP Statistical Processing Software (SAS, Cary, NC, USA) or custom-written Matlab code (Mathworks, Natick, MA).

## RESULTS

### Normally-reared and song-naïve females engage in similar pair-bonding behaviors

Females reared with exposure to adult song (‘normally-reared’) and those reared with mothers and siblings but no adult song exposure (‘song-naïve’) were housed either with a male (‘mated’ females) or with another female (‘unmated’ females) for a period of two weeks (see Methods, Figure 1A). We recorded the display of a number of individual and interactive behaviors of male-female pairs, including courtship (female-directed or FD) and non-courtship (undirected or UD) singing, affiliative behaviors (clumping, allopreening), aggressive behaviors (bill fencing, pecking), and nesting behaviors (building, arranging, incubating), immediately after introduction (Week 0), and after one (Week 1) and two (Week 2) weeks of cohabitation. We found that male-female pairs exhibited a progression of behaviors over the two-week cohabitation period (Figure 1B). In general, over the course of two weeks, there were changes in the expression of courtship (FD singing F_2,42_= 14.86, p<0.001), affiliative (preening F_2,42_=4.38, p=0.0187), aggressive (fencing F_2,42_ = 8.14,p=0.001), and nesting (F_2,42_=13.82, p<0.0001) behaviors. Pairs displayed courtship behaviors and aggression during the first introduction; these behaviors significantly decreased by week 1 and remained low thereafter (week 0 > week 1, 2, p<0.05 for FD singing, fencing; Figure 1C). In contrast, other affiliative behaviors and nest activities (building, arranging, incubating) increased after the first introduction. For example, allopreening increased in week 1 and remained high in week 2 (week 1 > week 0 p=0.0253; week 2 > week 0 p=0.0564), while pairs increased the time spent nest building and arranging in week 1 and increased even more in week 2 (week 1,2 > week 0 p<0.005 Figure 1C). Clumping remained relatively stable, showing a slight but not significant increase between the introduction and the end of week 1 (F_2,42_=1.31, p=0.2818; Figure 1C). Pecking remained low across the three weeks.

During the two week period of cohabitation, both normally-reared and song-naïve pairs displayed strikingly similar patterns of social behavior (Figure 1B,C). Indeed, there were no significant effects of rearing or rearing by time interactions on either affiliative or aggressive behaviors (p>0.40 for courtship singing, allopreening, clumping, and aggressive behaviors). While we generally observed that song-naïve females were later in initiating nesting activity, this difference was not significant (F_1,21_=3.65, p=0.0697). These data indicate that, even without exposure to a male or to male song during development, female zebra finches engage in remarkably similar patterns of social behavior during mating and pair bonding.

While the interactive behaviors were generally similar between the rearing conditions, one difference between normally-reared and song-naïve females was in the time spent in the nest. In particular, we found that by the end of the second week, normally-reared females were spending, on average, 44% (range 0-79%) of their time in the nest, significantly more than the 14% (range 0-61%) of time spent by song-naïve females (Figure 1C; rearing x time interaction F_2,42_= 3.54;p=0.0379; post-hoc comparisons normally-reared week 2 vs. song-naïve week 2 p=0.0128). Moreover, by the end of week 2, 8 of 11 normally-reared females spent more than 5% of their time in the nest, while only 4 of 12 song-naïve females were in the nest for more than 5% of the time (Figure 1C). Normally-reared females spent more time interacting than song-naive females during week 1 and week 2 due to this difference in nesting time (Figure 1B,C).

Thus, with the exception of nesting behavior, social interactions were similar between the rearing conditions.

### Pair bonding leads to mate preference formation in both normally-reared and song-naïve females

Following two-weeks of cohabitation, we tested females for their song preferences using an active choice assay in which females pulled strings to trigger song playback (see Methods) [33]. In general, both normally-reared and song-naïve females triggered song playback in the assay at similar rates (F_1,54_= 0.0267, p=0.8709) and this was not affected by mating experience (mating F_1,54_= 0.0150, p=0.9030; mating x experience interaction F_1,54_= 1.5069, p=0.2249). However, mating experience significantly influenced the direction and strength of females’ biases for a male’s song. We quantified the strength of preference between the two songs, by calculating a normalized preference index from the number of pulls in a test for each female. We found that mated females preferred the song of their mate over the song of an unfamiliar male while unmated females did not show similar bias toward the familiar song (Figure 2A; effect of experience F_1,54_=16.91, p<0.0001). Moreover, while recent social experience significantly affected preferences for the song of a familiar over an unfamiliar male, preferences were not significantly different between females from the two rearing conditions (Figure 2A; F_1,54_=0.87, p=0.3564). In particular, we found that the distribution of preferences was significantly greater than 0.5, indicating a preference for the mate’s song, for both mated normally-reared females (t=3.36, p=0.0073) and mated song-naïve females (t=2.95, p=0.0122; Figure 2A). Finally, we used the confidence intervals around the bootstrapped mean to determine when an individual displayed a significant preference for the mate’s song, the unfamiliar song, or no preference. We found that mated females from both rearing conditions showed significant preferences for the mate’s song on a greater proportion of tests than unmated females (Figure 2B). In contrast, we found that not only did females passively exposed to a male’s song (unmated females) not show consistent bias for the familiar song, but unmated normally-reared females showed a significant preference for the unfamiliar male’s song (t=-2.38, p=0.0276; Figure 2A). Compared to females in the other mating and rearing conditions, unmated song-naive females had the highest proportion of tests on which they showed no significant preferences (Figure 2B). Taken together, these data demonstrate that social experience with a male is similarly effective at driving learning and plasticity of song preferences in song-naïve and normally-reared females.

**Figure 2.**
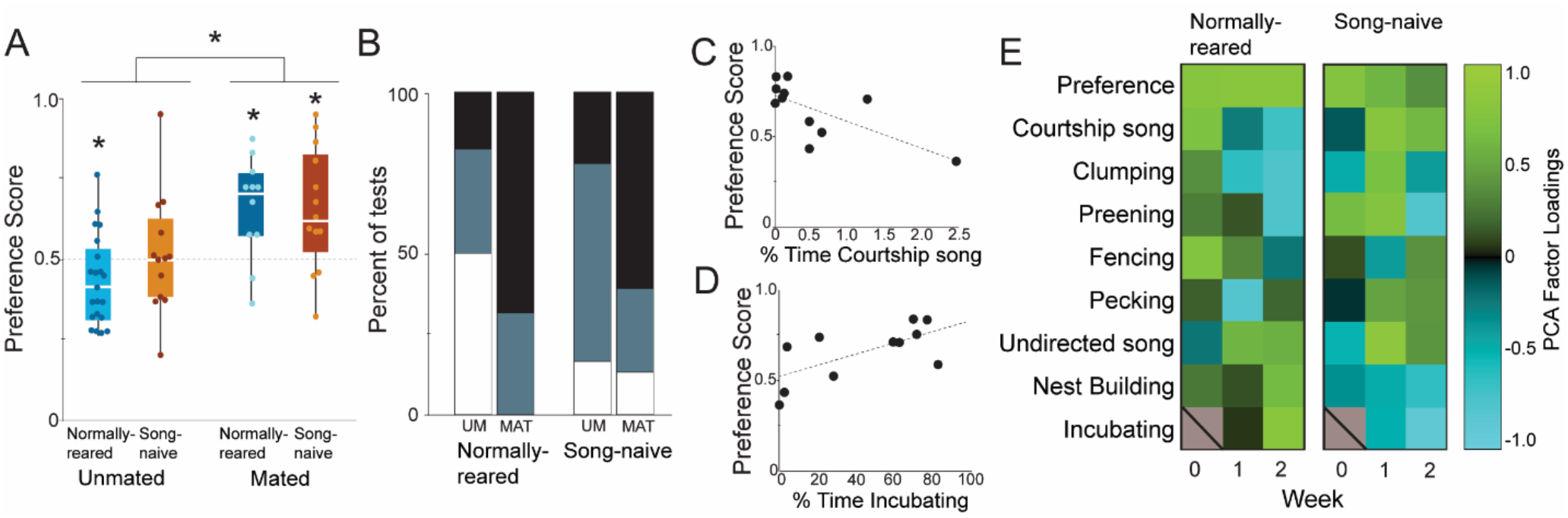
Mating influences learning of a mate’s song preference. A) Box and whisker plots of mated and unmated normally-reared (blues) and song-naïve (reds) females. Mated females from both rearing conditions show significant preferences for the song of a mate. In contrast, unmated normally-reared females prefer an unfamiliar song. Preference score is the bootstrapped preference value where 1 is a preference for the mate’s song, 0 is a preference for an unfamiliar song, and 0.5 indicates no preference. B) Percent of tests in which mated (MAT) and unmated (UM) females exhibited either a significant preference for the mate’s song (black), a significant preference for the unfamiliar song (white) or no significant preference (gray). Mated females from both rearing conditions prefer the mate’s song more frequently than unmated females. Unmated, normally-reared females show stronger preferences for the unfamiliar song. In normally-reared females, lower levels of courtship singing in week 2 (C) and (D) greater time spent incubating in week 2 are correlated with higher preference scores (see A for definition). E) Principal components analysis of preference and interactive behaviors for normally-reared (left) and song-naïve (right) females for weeks 0, 1, and 2. Plotted are heat maps of the normalized loading scores of the principal components with the strongest loading of preference. Normally-reared females show positive loadings of courtship and affiliative behaviors in week 0 which transition to negative loadings of courtship and affiliative behaviors and positive loadings of nesting behaviors in week 2. Song-naïve females show a distinctly different pattern of loadings.

Moreover, hearing the vocalizations of a male without physical or visual interactions is not sufficient to drive song preference towards a familiar song and may lead to preference for a novel song.

### Social interactions during pair-bonding influence females’ preferences for their mate’s song

Both normally-reared and song-naïve females show similar overall behaviors during two weeks of cohabitation and both show significant preferences for their mate’s song. We investigated whether the strength of preference in normally-reared and song-naïve females was predicted by the same, specific pair-bonding behaviors. We first looked for correlations between the preference for mate’s song and each of the pair bonding behaviors over the three time points. For normally-reared females, stronger mate’s song preferences were correlated with lower levels of courtship singing (Figure 2B; r=-0.6927, p=0.0181) and greater time spent incubating eggs (Figure 2C; r=0.6596, p=0.0273) on week 2. In contrast, for song-naïve females there were no significant correlations between mate’s song preference and any pair bonding measures.

We also used a principal components analysis (PCA) to probe whether particular linear combinations of pair bonding behaviors were differentially associated with mate’s song preference in the two rearing conditions. Among normally-reared females, stronger preferences for the mate’s song were associated with species-typical progression in pair-bonding behaviors (Figure 2D). In particular, preference loaded positively on PCs with greater courtship and affiliative behaviors at the beginning of pairing and with greater nesting and incubation behaviors in subsequent weeks. Specifically, in week 0, the PC with the highest preference loading also had substantial positive loading of courtship and associative behaviors (clumping, preening, courtship song) and negative loading of nesting (Figure 2D). Over the course of weeks 1 and 2, this pattern reversed such that by week 2 the PC with the highest loading of preference also had high, positive loadings of nesting, incubation, and undirected song, and negative loadings for courtship song and affiliative behaviors. The pattern in song-naïve females was markedly different. The PC with the highest preference loading in week 0 had positive loading of preening, and weak or negative loading of all other behaviors. By week 2, positive loading of preference was associated with singing and aggressive behaviors, and weak or negative loading of affiliative and nesting behaviors. Taken together, these data indicate that while normally-reared and song-naïve females show similar progressions of pair bonding behaviors over time and similar preference formation for the mate’s song, there are substantial differences in the relationship between preference and pair bonding behaviors between the rearing conditions.

Previous work has shown that mated females show strong preferences not only for the song of a mate over an unfamiliar male, but also for their mate’s courtship song over his non-courtship song [21]. Thus, we also tested the degree to which rearing and mating experience influenced the preference for the mate’s courtship song. We found that mating experience significantly affected the preference for courtship over non-courtship song (Figure 3A; F_1,38_= 5.4076, p=0.0255). However, this effect was driven primarily by the responses of mated, normally-reared females, who showed significant preferences for the mate’s courtship vs. non-courtship song (Figure 3A; t=3.12, p=0.0169). (Figure 3B). In contrast, the preferences of mated song-naïve females were not significantly different from 0.5 (t=1.13, p=0.2965). Neither group of unmated females (normally-reared or song-naïve) had preferences that differed significantly from 0.5 (p>0.40 for both). Finally, in addition to showing a bias towards the mate’s courtship song, mated, normally-reared females never showed significant preferences for the mate’s non-courtship song (Figure 3B). In contrast, unmated normally-reared females and song-naive females from both conditions showed significant preferences for the non-courtship tests on 25-55% of tests.

**Figure 3.**
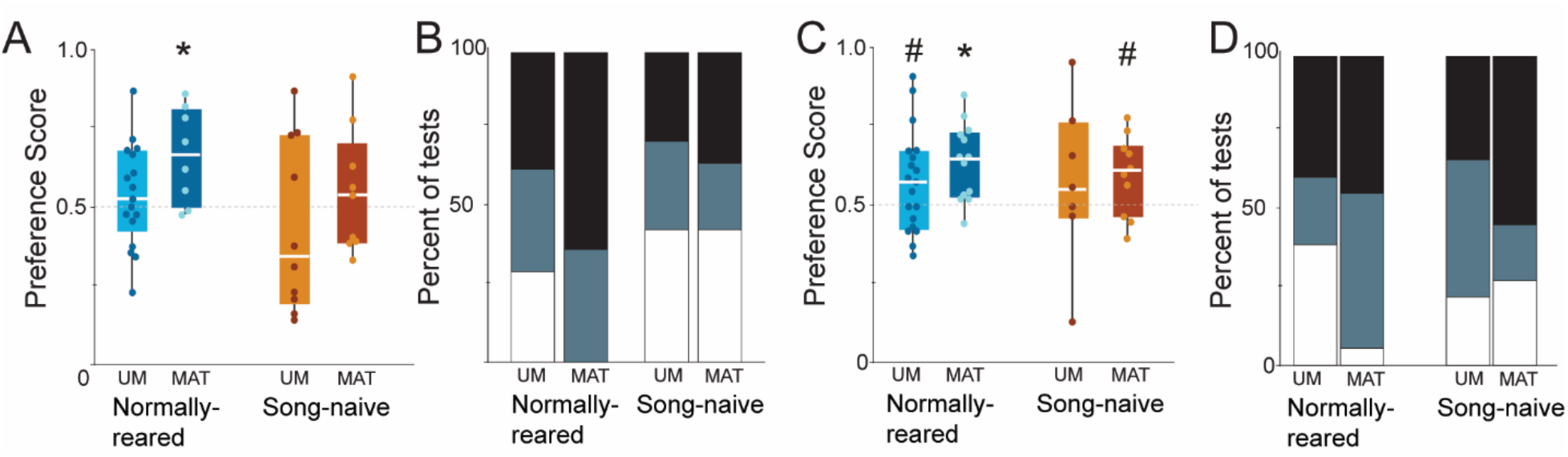
Effects of rearing condition and mating experience on preferences for courtship song. A) Only mated (MAT), normally-reared females have preference scores significantly greater than 0.5 for the courtship songs of their mate. Box and whisker plots of the preference score for mated and unmated normally-reared and song-naïve females tested on courtship vs. non-courtship songs of their mate. Preference score is the bootstrapped preference value where 1 is a preference for courtship song, 0 is a preference for non-courtship song, and 0.5 indicates no preference. Median is shown in white, points are responses of individual females. (B) Percent of tests on which females showed significant preference for the mate’s courtship song (black), non-courtship song (white) or no preference (gray). Mated, normally-reared females more frequently prefer the mate’s courtship song compared to all other groups. (C) Box and whisker plots of the preference score for mated and unmated normally-reared and song-naïve females tested on unfamiliar courtship vs. non-courtship songs. Preference scores of mated, normally-reared females are significantly greater than 0.5, indicating preferences for the courtship song. Both unmated, normally-reared females and mated song-naïve females show a trend in the same direction. (D) Percent of tests on which females showed significant preference for courtship song (black), non-courtship song (white), or no preference (gray). Mated, normally-reared females are less likely to prefer the non-courtship song compared to the other groups. Symbols indicate groups that are significantly different from chance (0.50). *p<0.05, #p<0.10.

### Developmental song exposure and recent social experience interact to affect courtship song preference

Finally, we sought to determine whether mating experience can influence preferences for songs other than the song of the mate. Previous work has found developmental song exposure shapes species-typical preferences for unfamiliar songs. In particular, unlike normally-reared females, song-naïve females do not show consistent preferences for courtship song versus non-courtship song from an unfamiliar male. Here, we investigated whether pair-bonding could also lead to consistent courtship song preferences in song-naïve females toward unfamiliar courtship vs. non-courtship songs. There were no significant effects of rearing, mating experience, or an interaction on the bootstrapped preference values (p>0.50 for all comparisons). However, the degree of deviation from 0.5 did vary across groups. Normally-reared, mated females were significantly above 0.5, indicating preferences for unfamiliar courtship songs over unfamiliar non-courtship songs (Figure 3B; t=4.68, p=0.0016). Unmated normally-reared females, and mated song-naïve females tended towards a deviation above 0.5, however, this was not statistically significant for either group in two-tailed tests (Figure 3B; unmated, normally-reared t=1.78, p=0.0960; mated, song-naïve t=1.97, p=0.0850). Unmated, song-naïve females showed the greatest variation across individuals and did not deviate from 0.5 (p>0.40). Finally, mated, normally-reared females rarely preferred non-courtship songs, while unmated, normally-reared females and song-naive females from both mating conditions tended to show a preference for non-courtship songs on 10-25% of tests (Figure 3D).

## DISCUSSION

Variation in social and sensory experience, both during development and in adulthood, has the potential to shift the trajectory of adult social interactions and relationships. In particular, adult experiences can have dramatic effects on sensory plasticity and perception [6,40,41]. However, the degree to which developmental organization of sensory systems during a critical period may contribute to or limit the impact of adult social or sensory experience is less clear. Here, we investigated the impact of developmental exposure to species-typical communication signals (songs) on adult social interactions and learned song preferences in zebra finches. We found that female zebra finches that were isolated from male song during development (‘song-naïve’ females) displayed similar social and mating interactions to birds reared with species-typical song exposure (“normally-reared” females). Despite the lack of prior experience with any male song, song-naive females were indistinguishable from normally-reared females in their preference learning of the mate’s song following two weeks of co-housing with the mate. In contrast, passive exposure to the male’s song was not sufficient to induce song preference. Mating experience was not sufficient, however, to fully rescue other species-typical auditory preferences in song-naive females, such as preference for the mate’s courtship song over his non-courtship song. Moreover, though we observed similar patterns of pair bonding behaviors in all male-female pairs, the relationships between pair bonding behaviors and the strength of preference differed between normally-reared and song-naïve females. Taken together, these data indicate that although song-naïve females are able to form pair bonds and preferences for their mate’s song following two weeks of cohabitation, the lack of song exposure during development has long lasting effects on auditory processing and social interactions that are not ameliorated by adult social and mating experience.

Early life rearing conditions can affect the expression of adult prosocial behaviors [42]. For example, male and female prairie voles reared with single mothers show delayed partner preference formation compared to voles reared with male/female breeding pairs [42]. In our study, rearing condition did not significantly affect the overall progression of social and pair bonding behaviors. Both normally-reared and song-naïve birds moved from courtship and affiliative behaviors during the first introduction, to nesting and incubation behaviors by the end of the second week. Moreover, both song-naïve and normally-reared females showed strong preferences for the song of their mate after two weeks of cohabitation. The remarkable consistency in the progression of pair bonding behaviors could result because behavioral transitions in pair bonding are not dependent on early rearing experience or song exposure in development, or, alternatively, because song-naïve females could follow the lead of their male mate, all of whom were normally-reared in our study. The main difference due to rearing was that fewer song-naïve females were sitting in nests and incubating eggs by the end of the second week compared to their normally-reared counterparts. One possibility is that reproductive or neuroendocrine physiology associated with breeding may have been delayed in song-naïve females due to the lack of exposure to song either during development or in the weeks leading up to pairing. Hearing song can stimulate the reproductive axis and increase egg-laying in zebra finches [43]. Thus, whether this difference in incubation is due to the lack of song stimulation immediately prior to pairing versus an effect of developmental influences requires further investigation.

While there was similarity in both the progression of pair bonding behavior and the preference for the mate’s song between the two rearing conditions, the relationships between preference and pair bonding behaviors were not similar. In normally-reared females, preference for the mate’s song was correlated with a species-typical progression of mating behaviors over the course of pair bonding, with stronger preferences correlating with courtship and affiliative behaviors during the first introduction and with nesting and incubation behaviors by the end of the second week. In song-naïve females, despite typical mating and social interactions overall, preference for the mate’s song was correlated with a different progression of mating behaviors over the two-week period. Together, these data raise the possibility that while pair bonding and preference appear similar between the rearing conditions, there may be different aspects of social and mating interactions that drive preference learning and plasticity in song-naïve females.

Hearing song during development may be important for both learning a tutor song or template, and for shaping auditory perception more generally. Passive tutoring of both male and female finches between 25-35 days can lead to preferences for the tutor song, and, in males, to tutor song copying [31,44]. Rearing female zebra finches without exposure to song not only leads to a lack of preference for a tutor song, but to diminished auditory perceptual abilities and aberrant song preferences compared to normally-reared females [26,45]. While song-naïve zebra finches can distinguish between species with acoustically distinct songs (e.g. zebra finch song vs. canary song), they show diminished absolute and relative pitch discrimination and inconsistent preferences for female-directed courtship song relative to non-courtship song [22,26]. Thus, developmental song exposure appears to be necessary for females to perform fine-tuned discrimination and perception. We hypothesized that if these deficits stem from the lack of a song ‘template’, mating could promote song memorization and auditory plasticity, leading to both a learned preference for the mate’s song and, potentially, to an amelioration of species-typical courtship song preferences. We found that although there was a trend toward stronger preferences for the courtship song, mated song-naïve females still preferred the non-courtship song more frequently than mated normally-reared females. While further investigation into the perceptual abilities of mated, song-naïve females are warranted, these data hint that mating is not sufficient to fully overcome the lack of exposure to song during development.

While mated females exhibited strong preferences for the familiar, mate’s song, unmated, normally-reared females showed significant preferences for a novel song over a familiar song. Previous work has found that male zebra finches show behavioral habituation to repeated song playback [46,47]. In contrast, female responses to passive song exposure appear dependent on their preference for the song and not the familiarity or novelty per se (Woolley & Barr, unpub data). This has led to the question of whether females form auditory memories of songs they are passively exposed to. The current finding that unmated, normally-reared females show a preference for the novel song indicates that females may indeed attend to songs they are repeatedly exposed to, and that they may track not only familiarity, but other social and acoustic cues. In particular, the lower preference for the familiar song indicates that females are sensitive not just to song acoustics, but also to the social context or dynamics in which the song is performed. In a diversity of species, females can assess a male’s fitness or availability by eavesdropping on his interactions with other conspecifics [48,49], and this information can influence reproductive decisions [50]. Our data indicate that female zebra finches may also be sensitive to the mating status and other social information conveyed in the vocal communication between a pair.

In adult animals, auditory plasticity can be induced through neuromodulatory effects on the auditory cortex. For example, pairing playback of a tone with stimulation of acetylcholine, or dopamine inputs to the auditory cortex can lead to expansion of the tonotopic representation of the tone frequency [51–53]. In more naturalistic contexts, recent studies on maternal behavior have found that oxytocin in the auditory cortex of new mothers enables neural plasticity necessary for changing the valence of pup calls [54]. In female zebra finches, pairing dopaminergic stimulation of the secondary auditory cortex with passive playback of a less preferred song can lead to a preference for a previously less preferred song [33]. Moreover, neurons in the caudal VTA are more active in response to preferred compared to less preferred songs [24]. We hypothesize that physical and audiovisual interactions with a mate can lead to enhanced activity of midbrain dopaminergic neurons. Neurons in the cVTA project to sensory regions, such as the NCM, which is important for song memory and shows differential responses to the song of a mate versus an unfamiliar male [21,33,55,56]. Rearing females without song exposure leads to both aberrant responses in the NCM and the VTA [22,24], and this may underlie differences in correlations between preference and social behavior.

Further investigation of these neural and behavioral connections, in particular the degree to which differences in sensory tuning drive changes in the VTA of song-naïve females, will lend new insight into the neuromodulatory mechanisms of life-long adaptive plasticity in sensory perception.

## ACKNOWLEDGEMENTS

We would like to thank Jon T. Sakata, Logan James, Nancy Chen, Isabella Catalano, and Helena Barr for helpful discussions and comments on the manuscript. In addition, we would like to thank Ani Muradyan, Taylor Morganstein, Therese Koch, Lily Xu, Helen Lai, Jill Belanger, India Blaisdell, Sabrina Hennecke for help with preference testing and video coding, and Emma Hudgins and Lee Wall for assistance with coding. This work was funded by the Natural Sciences and Engineering Research Council of Canada (NSERC) and Fonds de Recherche du Québec – Nature et technologies (FQRNT) to SCW, and a Center for Research in Brain, Language, and Music graduate stipend award and Lloyd Carr-Harris fellowship to EMW.

## REFERENCES

1. Bellono NW, Leitch DB, Julius D. 2018 Molecular tuning of electroreception in sharks and skates. Nature 558, 122–126. (doi:10.1038/s41586-018-0160-9)

2. Endler JA. 1992 Signals, Signal Conditions, and the Direction of Evolution. Am. Nat. 139, S125–S153. (doi:10.1086/285308)

3. Heimonen K, Immonen E-V, Frolov RV, Salmela I, Juusola M, Vähäsöyrinki M, Weckström M. 2012 Signal coding in cockroach photoreceptors is tuned to dim environments. J. Neurophysiol. 108, 2641–2652. (doi:10.1152/jn.00588.2012)

4. Grubb MS, Thompson ID. 2004 The influence of early experience on the development of sensory systems. Curr. Opin. Neurobiol. 14, 503–512. (doi:10.1016/j.conb.2004.06.006)

5. Morishita H, Hensch TK. 2008 Critical period revisited: impact on vision. Curr. Opin. Neurobiol. 18, 101–107. (doi:10.1016/j.conb.2008.05.009)

6. de Villers-Sidani E, Merzenich MM. 2011 Chapter 8 - Lifelong plasticity in the rat auditory cortex: Basic mechanisms and role of sensory experience. In Progress in Brain Research (eds AM Green, CE Chapman, JF Kalaska, F Lepore), pp. 119–131. Elsevier. (doi:10.1016/B978-0-444-53752-2.00009-6)

7. Sanes DH, Bao S. 2009 Tuning up the developing auditory CNS. Curr. Opin. Neurobiol. 19, 188–199. (doi:10.1016/j.conb.2009.05.014)

8. Bradbury JW, Vehrencamp SL. 2011 Principles of animal communication, 2nd ed. Sunderland, MA, US: Sinauer Associates.

9. Mock DW, Fujioka M. 1990 Monogamy and long-term pair bonding in vertebrates. Trends Ecol. Evol. 5, 39–43. (doi:10.1016/0169-5347(90)90045-F)

10. Sakata JT, Catalano I, Woolley SC. 2022 Mechanisms, development, and comparative perspectives on experience-dependent plasticity in social behavior. J. Exp. Zool. Part Ecol. Integr. Physiol. 337, 35–49. (doi:10.1002/jez.2539)

11. Young LJ, Wang Z. 2004 The neurobiology of pair bonding. Nat. Neurosci. 7, 1048–1054. (doi:10.1038/nn1327)

12. Carter CS, Keverne EB. 2002 4 - The Neurobiology of Social Affiliation and Pair Bonding. In Hormones, Brain and Behavior (eds DW Pfaff, AP Arnold, SE Fahrbach, AM Etgen, RT Rubin), pp. 299–337. San Diego: Academic Press. (doi:10.1016/B978-012532104-4/50006-8)

13. Johnson ZV, Young LJ. 2015 Neurobiological mechanisms of social attachment and pair bonding. Curr. Opin. Behav. Sci. 3, 38–44. (doi:10.1016/j.cobeha.2015.01.009)

14. Scribner JL et al. 2020 A neuronal signature for monogamous reunion. Proc. Natl. Acad. Sci. 117, 11076–11084. (doi:10.1073/pnas.1917287117)

15. Walum H, Young LJ. 2018 The neural mechanisms and circuitry of the pair bond. Nat. Rev. Neurosci. 19, 643–654. (doi:10.1038/s41583-018-0072-6)

16. Newman KS, Halpin ZT. 1988 Individual odours and mate recognition in the prairie vole, Microtus ochrogaster. Anim. Behav. 36, 1779–1787. (doi:10.1016/S0003-3472(88)80117-9)

17. Young KA, Gobrogge KL, Liu Y, Wang Z. 2011 The neurobiology of pair bonding: insights from a socially monogamous rodent. Front. Neuroendocrinol. 32, 53–69. (doi:10.1016/j.yfrne.2010.07.006)

18. Clayton NS. 1988 Song discrimination learning in zebra finches. Anim. Behav. 36, 1016–1024. (doi:10.1016/S0003-3472(88)80061-7)

19. Dong S, Clayton DF. 2009 Habituation in songbirds. Neurobiol. Learn. Mem. 92, 183–188. (doi:10.1016/j.nlm.2008.09.009)

20. Miller DB. 1979 Long-term recognition of father’s song by female zebra finches. Nature 280, 389–391. (doi:10.1038/280389a0)

21. Woolley SC, Doupe AJ. 2008 Social Context–Induced Song Variation Affects Female Behavior and Gene Expression. PLoS Biol. 6, e62. (doi:10.1371/journal.pbio.0060062)

22. Chen Y, Clark O, Woolley SC. 2017 Courtship song preferences in female zebra finches are shaped by developmental auditory experience. Proc. Biol. Sci. 284, 20170054. (doi:10.1098/rspb.2017.0054)

23. Paul A, McLendon H, Rally V, Sakata JT, Woolley SC. 2021 Behavioral discrimination and time-series phenotyping of birdsong performance. PLOS Comput. Biol. 17, e1008820. (doi:10.1371/journal.pcbi.1008820)

24. Barr HJ, Woolley SC. 2018 Developmental auditory exposure shapes responses of catecholaminergic neurons to socially-modulated song. Sci. Rep. 8, 11717. (doi:10.1038/s41598-018-30039-y)

25. Hauber ME, Woolley SMN, Theunissen FE. 2007 Experience-dependence of neural responses to social versus isolate conspecific songs in the forebrain of female Zebra Finches. J. Ornithol. 148, 231–239. (doi:10.1007/s10336-007-0234-1)

26. Sturdy CB, Phillmore LS, Sartor JJ, Weisman RG. 2001 Reduced social contact causes auditory perceptual deficits in zebra finches, Taeniopygia guttata. Anim. Behav. 62, 1207–1218. (doi:10.1006/anbe.2001.1864)

27. Alger SJ, Juang C, Riters LV. 2011 Social affiliation relates to tyrosine hydroxylase immunolabeling in male and female zebra finches (Taeniopygia guttata). J. Chem. Neuroanat. 42, 45–55. (doi:10.1016/J.JCHEMNEU.2011.05.005)

28. Day NF et al. 2019 D2 dopamine receptor activation induces female preference for male song in the monogamous zebra finch. J. Exp. Biol. 222, jeb191510. (doi:10.1242/jeb.191510)

29. Miller DB. 1979 The acoustic basis of mate recognition by female Zebra finches (Taeniopygia guttata). Anim. Behav. 27, 376–380. (doi:10.1016/0003-3472(79)90172-6)

30. Schubloom HE, Woolley SC. 2015 Variation in Social Relationships relates to song preferences and ERG1 expression in a female songbird. Dev. Neurobiol., 1–36. (doi:10.1002/dneu.)

31. Riebel K. 2003 Developmental influences on auditory perception in female zebra finches - Is there a sensitive phase for song preference learning? Anim. Biol. 53, 73–87. (doi:10.1163/157075603769700304)

32. Tchernichovski O, Mitra PP, Lints T, Nottebohm F. 2001 Dynamics of the Vocal Imitation Process: How a Zebra Finch Learns Its Song. science 291, 2564–2569.

33. Barr HJ, Wall EM, Woolley SC. 2021 Dopamine in the songbird auditory cortex shapes auditory preference. Curr. Biol. 31, 4547–4559.e5. (doi:10.1016/j.cub.2021.08.005)

34. Tchernichovski O, Nottebohm F, Ho CE, Pesaran B, Mitra PP. 2000 A procedure for an automated measurement of song similarity. Anim. Behav. 59, 1167–1176. (doi:10.1006/anbe.1999.1416)

35. Campbell DLM, Hauber ME. 2010 Behavioural correlates of female zebra finch (Taeniopygia guttata) responses to multimodal species recognition cues. Ethol. Ecol. Evol. 22, 167–181. (doi:10.1080/03949371003707885)

36. Klatt JD, Goodson JL. 2013 Sex-specific activity and function of hypothalamic nonapeptide neurons during nest-building in zebra finches. Horm. Behav. 64, 818–824. (doi:10.1016/j.yhbeh.2013.10.001)

37. Silcox AP, Evans SM. 1982 Factors affecting the formation and maintenance of pair bonds in the zebra finch, Taeniopygia guttata. Anim. Behav. 30, 1237–1243. (doi:10.1016/S0003-3472(82)80216-9)

38. Zann RA. 1996 The Zebra Finch: A Synthesis of Field and Laboratory Studies. Oxford University Press.

39. Kao MH, Brainard MS. 2006 Lesions of an Avian Basal Ganglia Circuit Prevent Context-Dependent Changes to Song Variability. J. Neurophysiol. 96, 1441–1455. (doi:10.1152/jn.01138.2005)

40. Keuroghlian AS, Knudsen EI. 2007 Adaptive auditory plasticity in developing and adult animals. Prog. Neurobiol. 82, 109–121. (doi:10.1016/j.pneurobio.2007.03.005)

41. Voss P, Thomas ME, Cisneros-Franco JM, de Villers-Sidani É. 2017 Dynamic Brains and the Changing Rules of Neuroplasticity: Implications for Learning and Recovery. Front. Psychol. 8.

42. Ahern T, Young L. 2009 The impact of early life family structure on adult social attachment, alloparental behavior, and the neuropeptide systems regulating affiliative behaviors in the monogamous prairie vole (Microtus ochrogaster). Front. Behav. Neurosci. 3.

43. Nakagawa S, Waas JR. 2004 The effect of acoustic and visual priming stimuli on the reproductive behaviour of female zebra finches, Taeniopygia guttata. Acta Ethologica 7, 43–49. (doi:10.1007/s10211-004-0097-x)

44. Derégnaucourt S, Poirier C, Kant AV der, Linden AV der, Gahr M. 2013 Comparisons of different methods to train a young zebra finch (Taeniopygia guttata) to learn a song. J. Physiol.-Paris 107, 210–218. (doi:10.1016/J.JPHYSPARIS.2012.08.003)

45. Lauay C, Gerlach NM, Adkins-Regan E, Devoogd TJ. 2004 Female zebra finches require early song exposure to prefer high-quality song as adults. Anim. Behav. 68, 1249–1255. (doi:10.1016/j.anbehav.2003.12.025)

46. Dai JB, Chen Y, Sakata JT. 2018 EGR-1 Expression in Catecholamine-synthesizing Neurons Reflects Auditory Learning and Correlates with Responses in Auditory Processing Areas. Neuroscience 379, 415–427. (doi:10.1016/j.neuroscience.2018.03.032)

47. Stripling R, Kruse AA, Clayton DF. 2001 Development of song responses in the zebra finch caudomedial neostriatum: Role of genomic and electrophysiological activities. J. Neurobiol. 48, 163–180. (doi:10.1002/neu.1049)

48. Cheney DL, Seyfarth RM. 2005 Social complexity and the information acquired during eavesdropping by primates and other animals. In Animal Communication Networks (ed PK McGregor), pp. 583–603. Cambridge University Press. (doi:10.1017/CBO9780511610363.030)

49. Otter K, McGregor PK, Terry AMR, Burford FRL, Peake TM, Dabelsteen T. 1999 Do female great tits (Parus major) assess males by eavesdropping? A field study using interactive song playback. Proc. R. Soc. Lond. B Biol. Sci. 266, 1305–1309. (doi:10.1098/rspb.1999.0779)

50. Mennill DJ, Ratcliffe LM, Boag PT. 2002 Female Eavesdropping on Male Song Contests in Songbirds. Science 296, 873–873. (doi:10.1126/science.296.5569.873)

51. Bakin JS, Weinberger NM. 1996 Induction of a physiological memory in the cerebral cortex by stimulation of the nucleus basalis. Proc. Natl. Acad. Sci. U. S. A. 93, 11219–11224.

52. Bao S, Chan VT, Merzenich MM. 2001 Cortical remodelling induced by activity of ventral tegmental dopamine neurons. Nature 412, 79–83. (doi:10.1038/35083586)

53. Kilgard MP, Merzenich MM. 1998 Cortical map reorganization enabled by nucleus basalis activity. Science 279, 1714–1718. (doi:10.1126/science.279.5357.1714)

54. Marlin BJ, Mitre M, D’amour JA, Chao MV, Froemke RC. 2015 Oxytocin Enables Maternal Behavior by Balancing Cortical Inhibition. Nature 520, 499–504. (doi:10.1038/nature14402)

55. Bolhuis JJ, Gahr M. 2006 Neural mechanisms of birdsong memory. Nat. Rev. Neurosci. 7, 347–357. (doi:10.1038/nrn1904)

56. Woolley SC, Woolley SMN. 2020 Integrating Form and Function in the Songbird Auditory Forebrain. In The Neuroethology of Birdsong (eds JT Sakata, SC Woolley, RR Fay, AN Popper), pp. 127–155. Cham: Springer International Publishing. (doi:10.1007/978-3-030-34683-6_5)

